# Annotation of piRNA source loci in the genome of non-model insects

**DOI:** 10.1101/2024.08.15.608080

**Authors:** Rebecca Halbach, Ronald P. van Rij

## Abstract

The PIWI-interacting RNA (piRNA) pathway plays a crucial role in the defense of metazoan genomes against parasitic transposable elements. The major source of piRNAs in the model organism *Drosophila melanogaster* are defective transposon copies located in piRNA clusters – genomic regions with a high piRNA density that are thought to serve as an immunological memory of past invasion by those elements. Different approaches have been used to annotate piRNA clusters in model organisms like flies, mice and rats, and software such as proTRAC or piClust are available for piRNA cluster annotation. However, these software often make assumptions based on current knowledge of piRNA clusters from (mostly vertebrate) model organisms, which do not necessarily hold true for non-model insects in which the piRNA pathway is less understood. Here we describe a simple piRNA cluster annotation approach that utilizes very little assumptions on the biology of the piRNA pathway. The pipeline has been validated on mosquito genomes but can be easily used for other non-model insect species as well.

## 1. Introduction

Transposable elements (TEs) are genetic parasites that make up a large fraction of eukaryotic genomes and are characterized by their ability to replicate and change position within their host genome [1]. The ubiquitous hazard of TEs requires potent defense mechanisms on the side of the host. In metazoans, the PIWI-interacting (pi)RNA pathway is the major defense against TEs [2]. piRNAs are small RNAs of approximately 23–32 nt that associate with PIWI clade proteins to repress TE expression and subsequent transposition in germline tissues [2]. The piRNA pathway is extensively studied in the fruit fly *Drosophila melanogaster*, where the majority of piRNAs are processed from precursors that are transcribed from large genomic regions called piRNA clusters, which are highly enriched in (defective or partial) TE sequences [3]. In germline cells, piRNAs are loaded onto the PIWI-protein Aubergine to degrade transposon transcripts in a positive feed-forward cycle termed the ping-pong amplification loop, involving Aubergine and another PIWI-protein, Argonaute3 [3, 4]. Alternatively, piRNAs can be loaded onto the eponymous Piwi protein to suppress TE expression by transcriptional silencing in both somatic and germline cells of the *Drosophila* ovary [5, 6].

There are two different types of piRNAs clusters in flies: dual-strand clusters that express piRNAs from both genomic strands and are expressed in germline cells, and uni-strand clusters which are biased towards piRNA expression from only one genomic strand and are the main source of piRNAs in somatic support cells in the gonads [3, 7]. Uni-strand clusters, such as the flamenco locus, are transcribed by RNA polymerase II (Pol II) as long piRNA precursors from a conventional upstream promoter and are subjected to capping, polyadenylation and splicing [8]. Dual-strand clusters, with *42AB* being the best-studied example, are marked by the repressive H3K9me3 heterochromatin modification [9] and lack conventional promoters [10]. Instead, the heterochromatin protein-1 paralog Rhino binds to H3K9me3 marks to allow for transcription within repressed chromatin. Along with its associated factor Deadlock, Rhino interacts with a homolog of the basal transcription factor TFIIA, called moonshiner, which recruits the Pol II pre-initiation complex and initiates internal transcription within the cluster region [11].

Identification of piRNA clusters is essential to uncover the biology and function of the piRNA pathway, but is especially challenging in understudied, non-model organisms. At the beginning of the piRNA era, clusters were defined in *Drosophila* and rodents by simply detecting regions in which the number of piRNAs exceeds a certain arbitrary threshold [3, 12]. However, threshold values suitable for piRNA cluster annotation in these animals are not necessarily transferable to other species, as they depend on the completeness or duplication level of (draft) reference genomes, biological factors such as relative piRNA expression, as well as technical variables such as the type of dataset and sequencing depth. To overcome these problems, software like proTRAC [13] and piClust [14] were developed. Though efficient and easy-to-use, these tools are optimized for vertebrate piRNA cluster annotation. As such, they rely on assumptions about nucleotide bias of piRNAs and strand bias of clusters that might not be known or might not hold true in other species, and they may therefore fail to annotate *bona fide* piRNA clusters. For example, these tools fail to identify noncanonical piRNA clusters like the tapiR locus in the yellow fever mosquito *Aedes aegypti*, which encodes piRNAs lacking a 1U bias that are essential for regulation of the maternal-to-zygotic transition [15].

The computational pipeline we describe here provides a simple tool for annotating piRNA clusters purely based on piRNA read density, while making as little assumptions about piRNAs and clusters as possible. With that, the pipeline can be especially suitable for species in which the piRNA pathway is not well characterized. The pipeline will 1) scan the genome for regions with a high piRNA coverage using non-overlapping 5-kb sliding windows, 2) merge these regions into larger clusters when in close proximity, 3) select clusters based on the presence of unambiguously mapping piRNAs, 4) determine the genomic start and end location of the clusters based on the location of the outermost piRNAs, and 5) eliminate very small clusters or clusters with very low piRNA density (Fig. 1). Since the annotation of piRNA clusters depends on arbitrary thresholds e.g., for piRNA coverage, density and other variables, our tool provides an option to extract basic summary statistics on the newly annotated clusters that can be easily used to explore and adjust cutoff threshold values to the studied species, type of dataset at hand, or the biological question of choice.

**Fig. 1.**
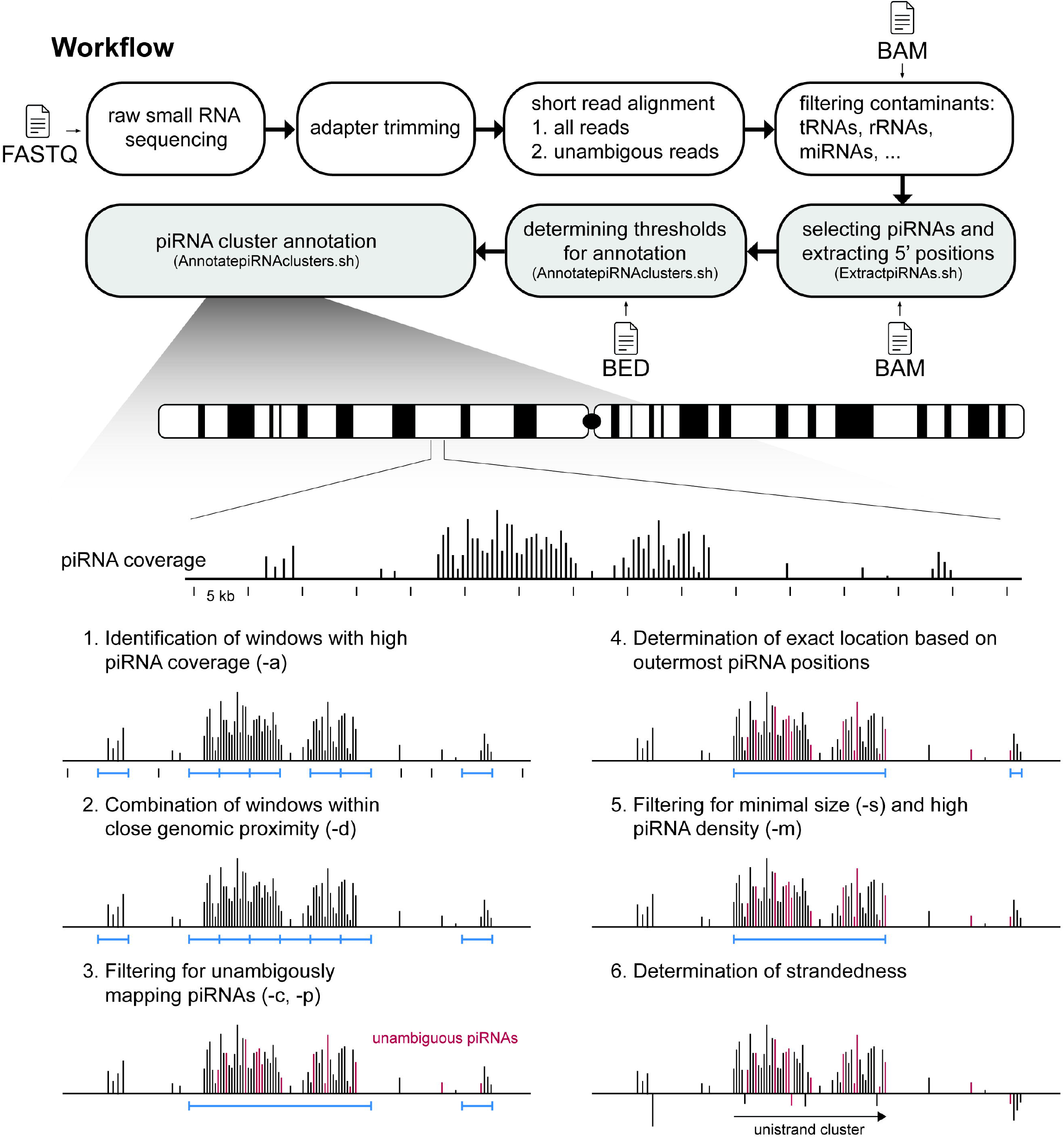
Schematic representation of principal steps of the piRNA cluster annotation pipeline. Small RNA-seq data are mapped to the genome and piRNA sized reads are selected (top). Then, (1) non-overlapping 5-kb sliding windows with high piRNA coverage are selected as putative piRNA clusters (blue horizontal lines), (2) these are merged into larger clusters when in close proximity, of which (3) clusters expressing unambiguously mapping piRNAs (red reads) are selected, (4) the genomic start and end locations of clusters are determined based on the location of the outermost piRNAs, (5) clusters are selected that exceed a minimal size and piRNA density, and finally (6) strandedness of the clusters in the final annotation is determined. Letters in brackets indicate the options that may be specified when executing the pipeline.

## 2. Materials

### 2.1. Required programs and scripts

The pipeline is based exclusively on freely available, open-source software that run on a Unix machine or cluster.

#### 2.1.1. Preliminary operations - Mapping of small RNA-seq libraries

- Cutadapt [16] (https://cutadapt.readthedocs.io/en/stable/)
- Bowtie [17] (https://bowtie-bio.sourceforge.net/index.shtml)
- Samtools [18] (https://www.htslib.org/)

#### 2.1.2. Filtering piRNA reads and annotation of piRNA clusters

- Bedtools [19] (https://bedtools.readthedocs.io/en/latest/)
- piRNA cluster annotation pipeline (https://github.com/vanrijlab/piRNAClusterAnnotation)

### 2.2. Computational power

The computational power required for executing the steps described below depends mostly on the size of the small RNA-seq library and the size of the reference genome. For mapping with Bowtie, processing memory (RAM) of at least 8 GB is recommended.

### 2.3. General considerations

1. The small RNA dataset: Total small RNA-seq datasets provide a comprehensive overview of piRNAs present in a given tissue and are suitable for the annotation of piRNA clusters. However, non-piRNA reads might contaminate the fraction of true piRNAs, and it can be difficult to distinguish between these two (see also below). Alternatively, small RNA-seq derived from PIWI immunoprecipitations that are enriched for genuine, PIWI-protein associated piRNA reads can be used as well. Yet, individual PIWI proteins might capture only a small subset of all piRNAs expressed in a given tissue, and, unless the binding repertoire of all expressed PIWI proteins can be acquired, will result in an incomplete piRNA cluster annotation.
2. The tissue to be analyzed: In most non-model arthropod species piRNAs are not restricted to the gonads but are expressed in somatic tissues as well [20]. Our pipeline allows for the annotation of piRNA clusters in different tissues separately using the same threshold values, since piRNA reads are normalized to the total piRNA pool instead of actual library depths. This enables the comparison of cluster annotations between different tissues, even if the amount of piRNAs relative to other classes of small RNAs differs significantly between these tissues. To get an overview of all piRNA clusters, the annotation can be performed with datasets from multiple tissues, and clusters can be merged afterwards to obtain a comprehensive list of piRNA clusters for a given species. Alternatively, cluster annotations from different tissues can be intersected to produce a list of core clusters that are ubiquitously expressed (e.g., [21]).
3. Removing contaminants: As indicated in 2.3.1, non-piRNA reads can contaminate the fraction of genuine piRNAs and result in false positive hits. If possible, these reads should be removed prior to running the cluster annotation pipeline. Most commonly, reads mapping to rRNA, tRNA, miRNAs, as well as other structured non-coding RNAs are computationally eliminated beforehand (see section 3.1.3). However, especially when using poorly annotated draft genome assemblies, annotation of such RNAs is often incomplete. In that case, it may be necessary to manually inspect and remove clusters that were wrongly defined by non-piRNA species.

## 3. Methods

This chapter mainly focuses on the annotation of piRNA clusters using mapped BAM files. Different alignment tools are available to map small RNA reads to a genome, and it is possible to use software of choice following procedures that work best for the organism of interest. However, for unexperienced users we also provide here the instructions to generate mapped BAM files from raw FASTQ files using the short read aligner Bowtie in the preliminary operation section (section 3.1.1)

### 3.1. Required files

The following files are required for mapping and pre-processing:

- Genome sequence in FASTA format.
- Raw small RNA sequencing read data in FASTQ format. 50 bp single-end sequencing is sufficient to capture small RNA sequences.
- Genome annotation file in GFF3 format.

The following files are required for using the piRNA cluster annotation pipeline:

- A BAM file containing small RNA reads mapped to the genome of interest. An example of how to trim adapter sequences with Cutadapt and align them with Bowtie is given in section 3.1.1.
- A tab-delimited chromosome file defining the chromosome lengths (section 3.1.3).

#### 3.1.1. Preliminary operation: Mapping of small RNA-seq data with Bowtie

This step requires Cutadapt and Bowtie to be installed. For installation guides, as well as more detailed information on the trimming and mapping procedures, we refer to the tools’ manuals. The steps for mapping small RNAs to a genome are: 1) trimming of adapter sequences from the reads and removing reads without adapter in the input FASTQ file (termed *input_file.fastq.gz* in the example below), 2) preparing bowtie index files for mapping, and 3) mapping trimmed reads to the genome. The last step has to be performed twice: once considering all reads and once retaining only unambiguously mapping reads.

For trimming and removing reads without adapters, run:

~~~
# Removing adapter sequence, and discarding reads that are much
shorter (<15nt) or longer (>35nt)than typical small RNAs
$ cutadapt -a [adapter_sequence] input_file.fastq.gz \
-m 15 -M 35 \
--discard-untrimmed |\
gzip > clipped_reads.fastq.gz
~~~

Bowtie requires an index of the reference genome for aligning the reads. This genome index file has to be created only once using the genome sequence in FASTA format (termed *genomesequence.fa in the example below*).

~~~
$ bowtie-build genomesequence.fa genome_index
~~~

Map the clipped small RNA reads to the genome, and create a sorted BAM file that contains all reads, regardless of whether they map once or multiple times to the genome. Note that it is important to randomly report only one alignment for ambiguously mapping reads to avoid inflation of the signal. Moreover, the numbers of mismatches allowed for mapping should be defined (Note 1).

~~~
$ bowtie genome_index clipped_reads.fastq.gz \
--best --strata \
-v [number_of_mismatches] -M 1 -S | \
# convert to BAM file
samtools view -Sb -F 4 - | \
# Sort BAM file
samtools sort - -o multimappers.bam
~~~

Create a second sorted BAM file that contains only reads that can be unambiguously assigned to a single genomic position:

~~~
$ bowtie genome_index clipped_reads.fastq.gz \
-v [number_of_mismatches] -m 1 -S | \
samtools view -Sb -F 4 - | \
samtools sort - -o uniquemappers.bam
~~~

#### 3.1.2. Optional: removal of rRNA, tRNA, miRNA reads and other contaminants

It can be useful to remove reads that are derived from rRNAs, tRNAs, (precursor) microRNAs, or other structured non-coding RNAs, unless they are interesting for the research question. For that, an annotation file in GFF3 format is required. The code below will search the GFF3 file for the terms rRNA, tRNA, and miRNA. Make sure that these terms are used as such in the file at hand, otherwise adjust the code accordingly. Extract genomic positions of contaminating non-coding RNAs:

~~~
$ awk ‘$3 ∼ /rRNA|tRNA|miRNA/’ annotationfile.gff > unwanted.gff
~~~

Then, remove all reads mapping to these genes with:

~~~
$ bedtools intersect -a multimappers.bam -b unwanted.gff -v >
multimappers_filtered.bam
~~~

#### 3.1.3. Creating a chromosome file

Chromosome sizes can be extracted with Samtools from a genome FASTA file with:

~~~
$ samtools faidx genomesequence.fa
$ cut -f1,2 genomesequence.fa.fai > chromfile.tab
~~~

Or, alternatively, from a BAM file (e.g., the file that was created in 3.1.1 with Bowtie) with:

~~~
$ samtools view -H multimappers.bam | \
grep @SQ | \
sed ‘s/@SQ\tSN:\|LN://g’ > chromfile.tab
~~~

#### 3.1.4. Size selection and extracting 5’ end positions of piRNAs

The piRNA cluster annotation pipeline will treat all reads that are provided in the input files as piRNAs and assumes that only the position of the 5’ end of the piRNA, not the entire read, is provided. It is therefore important to extract piRNA-sized reads and their 5’ end positions from the entirety of all small RNAs in the aligned BAM files before running the pipeline. For that, first clone the GitHub repository

~~~
$ git clone
https://github.com/vanrijlab/piRNAClusterAnnotation.git
$ cd ./piRNAClusterAnnotation
~~~

Make the script ExtractpiRNAs.sh executable with:

~~~
$ chmod +x ExtractpiRNAs.sh
~~~

Then, execute the code below for the two BAM files that were created in the previous steps and specify the minimal and maximum size of reads to be considered piRNAs (Note 2). For the second file, replace *multimappers.bam* with *uniquemappers.bam* (section 3.1.1 or 3.1.3):

~~~
$ bash ExtractpiRNAs.sh -i multimappers.bam \
-m [min_size_piRNA] -M [max_size_piRNA]\
-o [output_directory]
~~~

As output, this script will create a BED file with the input file’s name and the extension *.piRNAs.bed* containing the 5’ end position of reads only in the specified size range.

### 3.2. Annotation of piRNA clusters

#### 3.2.1. Running the cluster annotation pipeline

Make the script AnnotatepiRNAclusters.sh executable with:

~~~
$ chmod +x AnnotatepiRNAclusters.sh
~~~

The script requires three input files:

- A BED file containing the 5’ end positions of all piRNA reads (section 3.1.4)
- A BED file containing the 5’ end positions of unambiguously mapping piRNA reads (section 3.1.4)
- A chromosome file indicating the lengths of all chromosomes or contigs (section 3.1.3)

To execute the pipeline, run the script with:

~~~
$ bash AnnotatepiRNAclusters.sh -i multimappers.piRNAs.bed \
-u uniquemappers.piRNAs.bed \
-g chromfile.tab \
-a [min_piRNAs_per_window] \
-d [max_distance_windows] \
-c [min_unique_piRNAs] \
-p [min_unique_positions] \
-s [min_size_cluster] \
-m [min_piRNA_density] \
-o [output_directory]
~~~

Required command line options are:

- -i: a BED file containing the 5’ end positions of all piRNA reads including single and multimapping reads.
- -u: a BED file containing the 5’ end positions of only piRNA reads that map unambiguously
- -g: a tab-delimited chromosome file containing the size of all chromosomes (or contigs)

Optional command line arguments to specify threshold values are:

- -a: Minimal number of piRNAs that need to map to a window
- -d: Maximum distance between two windows to be merged
- -c: Minimal number of uniquely mapping piRNAs that need to map to a cluster
- -p: Minimal number of unique piRNA positions per cluster
- -s: Minimal size of a cluster in bp
- -m: Minimal piRNA density per kb of cluster
- -o: Folder in which results will be stored. If not specified, the current working directory will be used.
- -l: Run script in loop-mode (see section 3.2.2).
- -x: Run script in debugging-mode. In that case the script will print the code that is executed and will not delete temporary files.

Default values will be used if these threshold values are not specified by the user (Note 3).

The pipeline will create a log file that contains a short summary of the run and the results (*log.input_filename*), and a cluster file (*cluster.input_filename*) that contains all annotated piRNA clusters and basic mapping statistics:

| Column | Column name | Description |
| --- | --- | --- |
| 1 | Chrom | Chromosome or scaffold name |
| 2 | Start | Start coordinate of cluster (zero-based coordinate system) |
| 3 | End | End coordinate of cluster |
| 4 | Cluster_length | Length of the cluster [bp] |
| 5 | Total_piRNAs | Number of piRNA reads (not normalized) mapping to a cluster |
| 6 | piRNA_density | Number of piRNAs per million piRNA reads per kb mapping to a cluster |
| 7 | piRNAs_per_million | Number of piRNAs per million piRNAs reads mapping to a cluster |
| 8 | Unique_piRNAs_per_million | Number of unambiguously mapping piRNAs per million piRNA reads mapping to a cluster |
| 9 | Unique_piRNA_positions | Number of positions at which piRNAs map unambiguously |
| 10 | Perc_plus | Fraction of piRNAs in a cluster mapping to the positive strand [%] |
| 11 | Perc_min | Fraction of piRNAs in a cluster mapping to negative strand [%] |

High-confidence piRNA clusters are expected to have a high piRNA density, supported by a large fraction of unambiguously mapping piRNAs at multiple positions compared to total mapping piRNAs within the cluster, and they may be very large (e.g., the *flamenco* locus in *Drosophila melanogaster* is > 150 kb, and the top-10 most highly expressed piRNA clusters in *Aedes aegypti* range in size from 3 to > 300 kb). The annotated clusters should always be visually inspected, for example by using the online UCSC Genome Browser [22] or the desktop application Integrative Genomics Viewer (IGV) [23] (see Fig. 2A for examples).

**Fig. 2.**
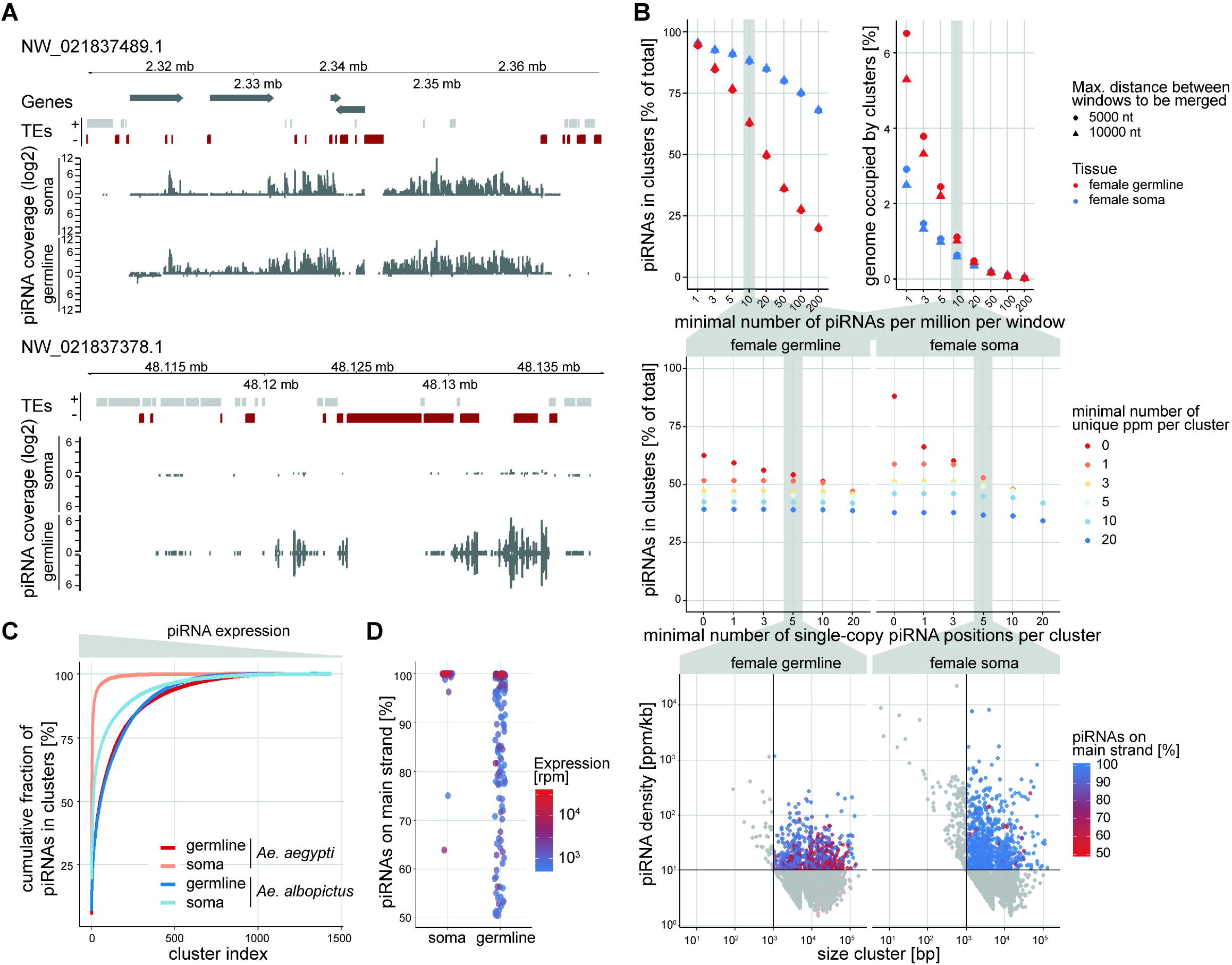
Examples of piRNA cluster annotation and characterization in Aedes mosquitoes. **A)** Two exemplary piRNA clusters in *Ae. albopictus*, expressed either from one genomic strand (uni-strand cluster, top panel), or both strands (dual-strand cluster, bottom panel). Gene annotation is indicated with dark gray arrows, and transposons are highlighted with gray and red boxes for positive or negative strand insertions, respectively. These piRNA clusters are large genomic regions with overall high piRNA coverage and density. **B)** Optimization strategy for piRNA cluster annotation in *Ae. albopictus*. Top panel: Influence of the minimal number of piRNAs required to map to a 5-kb window, and distance between windows (command line option -a and -d). The more stringent this cutoff is chosen, the fewer cluster will be annotated, likely resulting in many false negatives. Conversely, if a very low threshold is chosen, most piRNAs will be located in clusters, however, the annotation will include many false positives that might simply depict regions with background piRNA coverage. This might be different for different tissues, as indicated here for *Ae. albopictus* germline and somatic tissues. Middle panel: The effect of thresholds for the number of unique piRNAs (color-coded symbols; ppm, piRNAs per million mapped piRNAs) and unique piRNA positions (x-axis) on cluster annotation (command line options -c and -p), for germline (left panel) and somatic tissues (right panel). The more stringent these cutoffs are chosen, the more confident the annotation will be, however, the fewer clusters will be annotated. Bottom panel: Removal of very small loci, which may represent stand-alone transposon insertions but not piRNA clusters, and large loci with very low piRNA density that may reflect concatenated regions with background levels of piRNA expression but not *bona fide* clusters (command line options -s and -m). **C)** Cumulative fraction of all piRNAs located in annotated piRNA clusters in germline and somatic tissues of *Ae. aegypti* and *Ae. albopictus*. Cluster expression can be heavily biased towards a few large and highly expressed clusters, e.g., the three most highly expressed clusters account for more than half of all cluster-derived piRNAs in the soma of *Ae. aegypti*. **D)** Strand bias of piRNA clusters in somatic and germline tissues of Ae. albopictus. Only clusters with at least 500 piRNAs per million total reads (rpm) are included in the analysis. In this mosquito species, the most highly expressed clusters are all expressed from only one genomic strand, both in the soma and germline of the tissue. Panels A and B are adapted from ref. [28] (licensed under CC BY 4.0).

#### 3.2.2. Empirical evaluation of threshold values

Often, optimal threshold values for piRNA cluster annotation are not known. In that case it is possible to run the script in loop-mode with the additional option -l. By that, a third file (*stat.input_filename*) will be created that contains summary statistics not on individual clusters but on the entirety of annotated clusters when using different threshold values:

| T<br>o<br>t<br>a<br>l<br>s<br>t<br>a<br>t<br>i<br>s<br>t<br>i<br>c<br>s | Column | Column name | Description |
| --- | --- | --- | --- |
|  | 1 | Min_piRNAs | Cutoff values for all parameters that were used for the annotation |
|  | 2 | Max_dist |  |
|  | 3 | Min_unique_piRNAs |  |
|  | 4 | Min_unique_pos |  |
|  | 5 | Min_size |  |
|  | 6 | Min_density |  |
|  | 7 | No_clusters | Number of annotated clusters |
|  | 8 | Av_length | Average length of all clusters |
|  | 9 | Frac_genome | Fraction of the genome that is covered by piRNA clusters [%] |
|  | 10 | Frac_piRNAs_clusters | Fraction of piRNAs that map within annotated clusters [%] |

To test multiple values for different cutoffs, the script can be executed within a for loop, or within multiple nested for loops. For example, to test all combinations of values for the minimum number of piRNAs per window of 1, 5, 10, 50, or 100 piRNAs per million mapped piRNAs, and a minimal distance of 1000, 5000, or 10,000 bp for windows to still be merged, run the following command, while keeping all other values constant:

~~~
$ for min_piRNAs in 1 5 10 50 100; do
        for dist in 1000 5000 10000; do
              bash AnnotatepiRNAclusters.sh \
             -i multimappers.piRNAs.bed \
             -u uniquemappers.piRNAs.bed \
             -g chromfile.tab \
             -a “${min_piRNAs}” \
             -d “${dist}” \ [other options]
        done
done
~~~

Note that comparing many different conditions can take a very long time to run, depending on the available computational power.

The stat file can easily be used to visualize the performance of different cutoff values and can aid in the decision on the optimal threshold values to be used (see section 3.3).

### 3.3. Interpretation of the results

The annotation pipeline is based on the implicit assumption that piRNA clusters are large genomic regions with high piRNA density (two exemplary clusters are depicted in Fig. 2A) and that piRNA clusters are the main source of piRNAs. Under this assumption, thresholds for cluster annotation should be optimized to increase the fraction of piRNAs that map within annotated clusters, while keeping the clusters as compact as possible, e.g., only a small fraction of the total genome should be occupied by piRNA clusters. In Fig. 2B we present a data-driven decision for cluster prediction in the Asian tiger mosquito *Aedes albopictus* based on the summary statistics that are generated when running the pipeline in loop-mode (section 3.2.2). In general, decisions on threshold values will have to strike a fine balance between completeness and accuracy of the cluster prediction and will always be arbitrary to some extent. Moreover, the optimal conditions may differ depending on the aim of the study. For example, if a list of piRNA clusters as complete as possible is desired, more relaxed threshold values can be chosen, at the cost of annotating a larger part of the genome as piRNA clusters. However, if one is interested in identifying just a few, high-confidence piRNA clusters that can be used as basis for experiments, more strict cutoffs might be more suitable. This may especially be appropriate in species in which relatively few piRNA clusters express the majority of piRNA, such as in somatic tissues of *Aedes aegypti* (Fig. 2C).

Of note, while the assumption that piRNA precursors are transcribed from large genomic regions holds true for many vertebrate and invertebrate species [3, 12, 24-30], this assumption may be violated in some species. For instance, piRNAs in *C. elegans* are individually transcribed instead of processed from large cluster transcripts [31, 32]).

Annotation of piRNA clusters can be the first step in characterizing the piRNA pathway in non-model organisms. A non-exhaustive list of possible follow-up analyses is presented below:

1. Characterization of the repeat content of piRNA clusters compared with the rest of the genome. For example, piRNA clusters of *Drosophila melanogaster* are highly enriched for transposon sequences [3], while clusters in Aedes mosquitoes are not [27, 28].
2. Analysis of the strand bias of clusters. Uni-strand clusters express piRNAs mostly from one genomic strand, while dual-strand clusters produce piRNAs from both genomic strands (Fig. 2A, D). In fruit flies, the biology of these clusters is markedly different [8, 10, 11, 33, 34].
3. Comparison of piRNA cluster expression between different tissues. For example, clusters expressed in somatic or germline tissues might differ with regard to their transposon content, strand bias, or other interesting features (Fig. 2).

## 4. Notes

Note 1: The most stringent mapping procedure is to not allow any mismatches. However, it may be required to increase the number of mismatches allowed if small RNA-seq data from a different strain than the reference genome is used, or when the reference genome from a closely related species is used to map small RNAs.

Note 2: The default values for piRNA sizes are 25 (-m) to 30 nt (-M). However, the size range of piRNAs may differ between different species (see [20] for examples). It is therefore useful to determine the size profile of all small RNAs beforehand in order to estimate the size range of piRNAs. This can be quickly done by executing the command below and plotting the results in a histogram:

~~~
    $ samtools view multimappers.bam | cut -f10 | awk ‘{ print length }’
| sort | uniq -c > sizeprofile.tab
~~~

In general, piRNAs will be recognizable as a broad peak of larger sized RNAs, distinct from the siRNA and miRNA populations.

Note 3: The default values (-a min_piRNAs_per_window: 10; -d max_distance_windows: 5000; -c min_unique_piRNAs: 5; -p min_unique_positions: 5; -s min_size_cluster: 1000; -m min_piRNA_density: 10) are based on piRNA cluster annotations for *Aedes aegypti* and *Aedes albopictus* [27, 28]. The values were selected to maximize the fraction of piRNAs mapping to piRNA clusters, while minimizing the fraction of the genome covered by clusters (Fig. 2B). These threshold values will likely require adjustment for other species of interest.

## Acknowledgments

We thank Femke van Hout for testing the pipeline and critically reading the manuscript. This work was funded by the Human Frontiers Science Foundation (Research Grant number RGP0007/2017).

